# Computer Vision and Deep Learning for Environment-Adaptive Control of Robotic Lower-Limb Exoskeletons

**DOI:** 10.1101/2021.04.02.438126

**Authors:** Brokoslaw Laschowski, William McNally, Alexander Wong, John McPhee

**Affiliations:** Waterloo Artificial Intelligence Institute and Department of Systems Design Engineering at the University of Waterloo, Waterloo, ON, Canada; Waterloo Artificial Intelligence Institute and the Department of Systems Design Engineering at the University of Waterloo, Waterloo, ON, Canada

## Abstract

Robotic exoskeletons require human control and decision making to switch between different locomotion modes, which can be inconvenient and cognitively demanding. To support the development of automated locomotion mode recognition systems (i.e., high-level controllers), we designed an environment recognition system using computer vision and deep learning. We collected over 5.6 million images of indoor and outdoor real-world walking environments using a wearable camera system, of which ~923,000 images were annotated using a 12-class hierarchical labelling architecture (called the ExoNet database). We then trained and tested the EfficientNetB0 convolutional neural network, designed for efficiency using neural architecture search, to predict the different walking environments. Our environment recognition system achieved ~73% image classification accuracy. While these preliminary results benchmark Efficient-NetB0 on the ExoNet database, further research is needed to compare different image classification algorithms to develop an accurate and real-time environment-adaptive locomotion mode recognition system for robotic exoskeleton control.

## I. Introduction

The state-of-the-art in robotic exoskeleton control for human locomotion uses a hierarchical architecture, including high, mid, and low-level controllers [1]–[2]. The high-level controller is responsible for determining the user’s locomotor intent (e.g., climbing stairs, sitting down, or level-ground walking). The mid-level controller converts the locomotor activity from the high-level controller into mode-specific reference trajectories (i.e., the desired device state for each locomotion mode); this control level typically includes individual finite-state machines with discrete mechanical impedance parameters like stiffness and damping coefficients, which are manually tuned for different locomotor activities. The low-level controller calculates the error between the measured and desired device states and commands the robotic actuators to minimize the error via reference tracking and closed-loop feedback control [1]–[2].

High-level transitions between different locomotor activities remains a significant challenge. Most commercial exoskeletons require users to perform exaggerated movements or use hand controls to manually switch between locomotion modes [1]–[2]. Although accurate, such manual high-level control and decision making can be inconvenient and cognitively demanding. Researchers have been working on developing automated locomotion mode recognition systems using pattern recognition algorithms and data from wearable sensors like inertial measurement units (IMUs) and surface electromyography (EMG) [1]–[2]. Whereas mechanical and inertial sensors respond to the user’s movements, the electrical potentials of biological muscles, as recorded using surface EMG, precede movement initiation and thus could predict locomotion mode transitions with small prediction horizons. Several researchers have combined mechanical sensors with EMG for automated intent recognition [3]–[5]; such neuromuscular-mechanical data fusion has shown to improve the locomotion mode recognition accuracies and decision times compared to implementing either system individually. However, these measurements are user-dependent, and surface EMG require calibration and are susceptible to fatigue, motion artifacts, changes in electrode-skin conductivity, and crosstalk between muscles [1].

Information about the walking environment can supplement these automated locomotion mode recognition systems based on neuromuscular-mechanical data. Environment sensing and classification would precede modulation of the user’s muscle activations and/or walking biomechanics, therein allowing for more accurate and robust high-level transitions between different locomotor activities. Studies have shown that supplementing an automated high-level controller with terrain information improved the classification accuracies and decision times compared to excluding the environmental context [4]–[5]. Common wearables used for environment sensing include radar detectors [6], laser rangefinders [4]–[5], [7], RGB cameras [8]–[13], and 3D depth cameras [14]–[19] (Fig. 1).

**Fig. 1.**
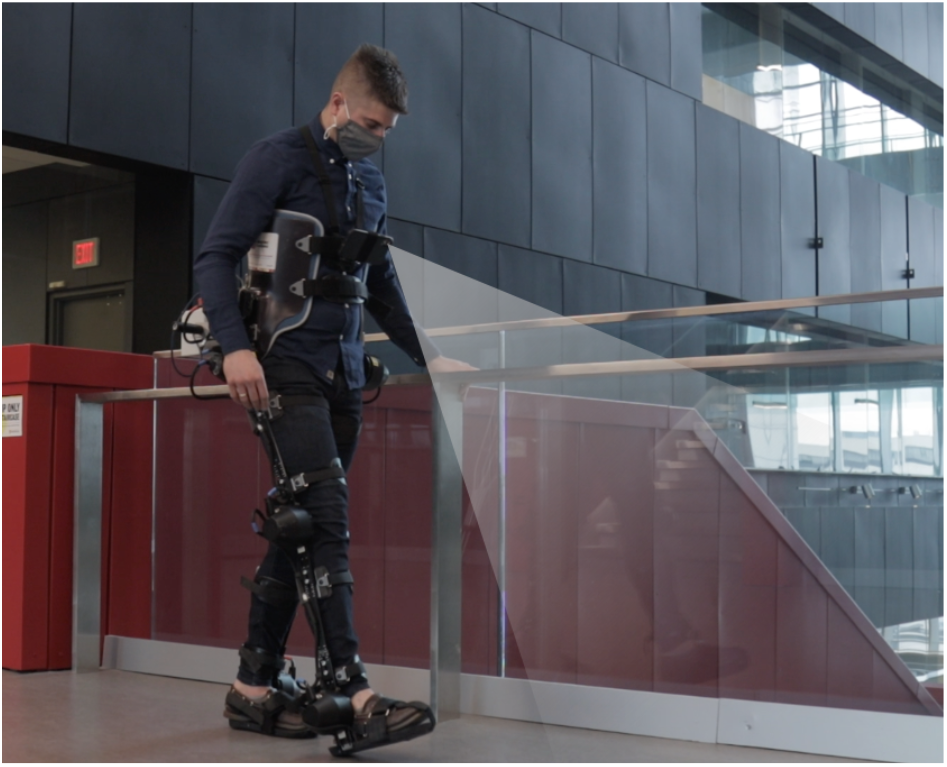
Photograph of the lead author wearing our robotic exoskeleton with environment sensing superimposed.

For classifying images of walking environments, researchers have used support vector machines [16]–[17] and convolutional neural networks (CNNs) [8], [10], [12]–[13], [18]–[19]. Although CNNs typically outperform support vector machines for image classification, deep learning requires significant and diverse training data to prevent overfitting and promote generalization. To date, researchers have each individually collected training data to develop their image classification algorithms. These repetitive measurements are time-consuming and inefficient, and individual private datasets have prevented comparisons between classification algorithms from different researchers [20]. We hypothesized that, by curating the largest and most diverse image dataset of walking environments, we could develop an environment recognition system using state-of-the-art convolutional neural networks to predict different real-world environments for robotic exoskeleton control; making the dataset open-source would also facilitate comparisons between next-generation environment recognition systems.

## II. Methods

### A. Experimental Dataset

One participant (without an exoskeleton) was instrumented with a wearable smartphone camera system (iPhone XS Max) (Fig. 2). Unlike limb-mounted systems [6]–[12], [16]–[17], [19], our chest-mounted camera can provide more stable video recording and allow users to wear pants and dresses without obstructing the field-of-view. The chest-mount height was ~1.3 m from the ground when the participant stood upright. The smartphone weighs ~0.21 kg and has an onboard re-chargeable lithium-ion battery, 512-GB of memory storage, and a 64-bit ARM-based integrated circuit (Apple A12 Bionic) with six-core CPU and four-core GPU; these hardware specifications can theoretically support onboard real-time inference. The relatively lightweight and unobtrusive nature of the wearable camera system allowed for unimpeded locomotion. Ethical review and approval were not required for this research in accordance with the University of Waterloo Office of Research Ethics.

**Fig. 2.**
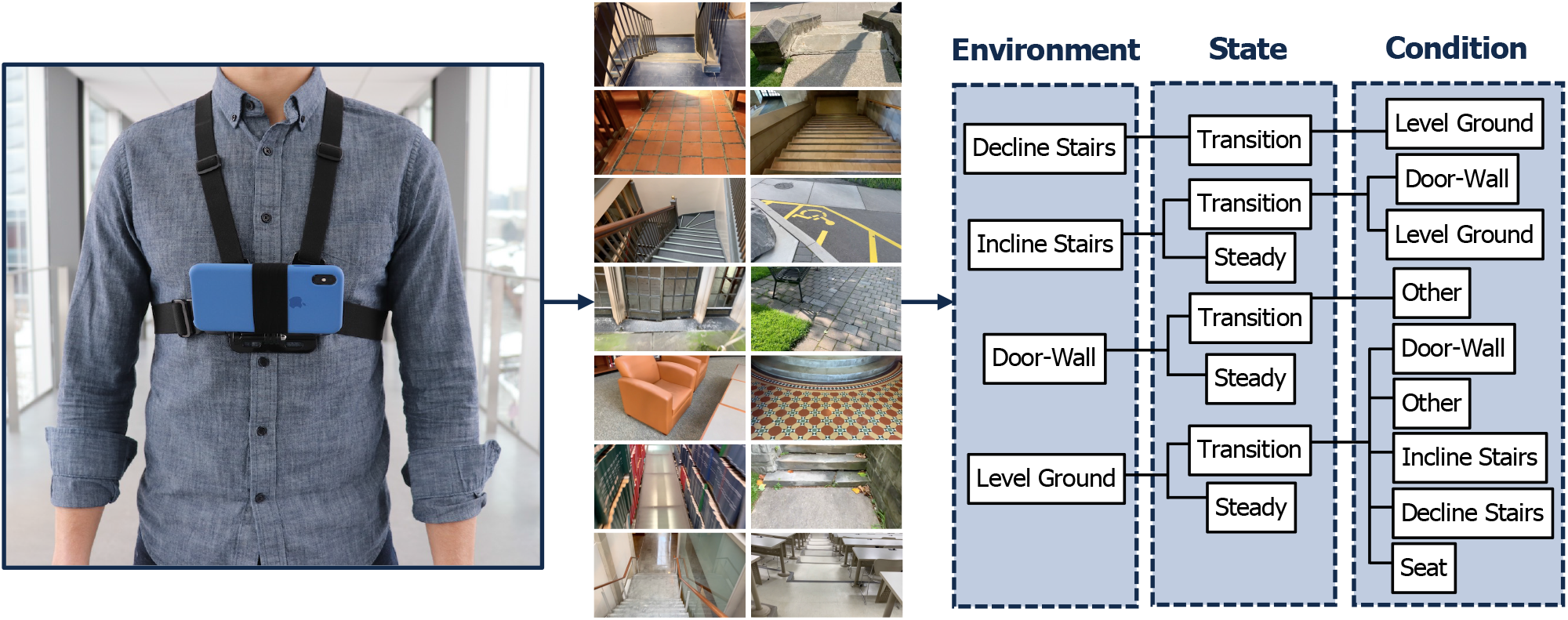
Development of the ExoNet database, including (left) photograph of our wearable camera system used for large-scale data collection; (middle) examples of the high-resolution RGB images of walking environments; and (right) schematic of the 12-class hierarchical labelling architecture [11].

Whereas previous studies have been limited to controlled indoor environments and/or prearranged walking circuits [3]–[9], [14]–[15], [18]–[19], our participant walked around unknown outdoor and indoor real-world environments while collecting images with occlusions, signal noise, and intraclass variations. We collected data at various times throughout the day to include different lighting conditions. The field-of-view was 1-5 m ahead of the participant. The camera’s pitch angle slightly differed between data collection sessions. Images were sampled at 30 Hz with 1280×720 resolution. We recorded over 52 hours of video, amounting to ~5.6 million images (Fig. 2). Data were collected throughout the summer, fall, and winter seasons to incorporate different weathered surfaces like snow, grass, and multicolored leaves. The image database, which we named ExoNet, was deposited in the IEEE DataPort repository and is now publicly available for download [11]. The file size of the uncompressed videos is ~140 GB.

Since there were minimal differences between consecutive images sampled at 30 Hz, we labelled the images at 5 frames/second. Approximately 923,000 images were manually annotated using a 12-class hierarchical labelling architecture (Fig. 2). The dataset included: 31,628 images of “incline stairs transition wall/door” (I-T-W); 11,040 images of “incline stairs transition level-ground” (I-T-L); 17,358 images of “incline stairs steady” (I-S); 28,677 images of “decline stairs transition level-ground” (D-T-L); 19,150 images of “wall/door transition other” (W-T-O); 36,710 images of “wall/door steady” (W-S); 379,199 images of “level-ground transition wall/door” (L-T-W); 153,263 images of “level-ground transition other” (L-T-O); 26,067 images of “level-ground transition incline stairs” (L-T-I); 22,607 images of “level-ground transition decline stairs” (L-T-D); 119,515 images of “level-ground transition seats” (L-T-E); and 77,576 images of “level-ground steady” (L-S). These class labels were chosen to encompass the different walking environments from the data collection. We included the *other* class to improve image classification performance when non-terrain related features like pedestrians, cars, and bicycles were observable.

Inspired by previous work [3]–[5], [8], our labelling architecture included both steady (S) and transition (T) states. A steady state describes an environment where an exoskeleton user would continue to perform the same locomotion mode (e.g., an image showing only level-ground terrain). In contrast, a transition state describes an environment where an exoskeleton high-level controller might switch between different locomotion modes (e.g., an image showing level-ground terrain and incline stairs). Manually labelling these transition states was relatively subjective. For example, an image showing level-ground terrain was labelled “level-ground transition incline stairs” (L-T-I) when an incline staircase was approximately within the field-of-view and forward-facing. Similar labelling was applied to transitions to other conditions.

### B. Convolutional Neural Network

We used TensorFlow 2.3 and the Keras functional API to train and test a convolutional neutral network for environment classification. During data preprocessing, the images were cropped to an aspect ratio of 1:1 and downsampled to 256×256 using bilinear interpolation. Random crops of 224×224 were used as inputs to the network; this method of data augmentation helped further increase the sample diversity. We used the EfficientNetB0 convolutional neural network developed by Google Research [21] for classification (Table 1). Unlike previous studies that used statistical pattern recognition or machine learning [14]–[17], deep learning models like Efficient-NetB0 can automatically and efficiently learn the optimal image features from the training data. The EfficientNetB0 architecture was designed using a multi-objective neural architecture search that optimized both classification accuracy and computational complexity [21]; these operational design features are important for real-time exoskeleton control. The final densely connected layer of the EfficientNetB0 architecture was modified by setting the number of output channels equal to the number of environment classes. Softmax was used to estimate the probability distribution (i.e., scores) for each environment. The network contained ~4 million parameters and ~391 million multiply-accumulate operations (MACs), which are representative of the architectural and computational complexities, respectively.

**Table 1.**
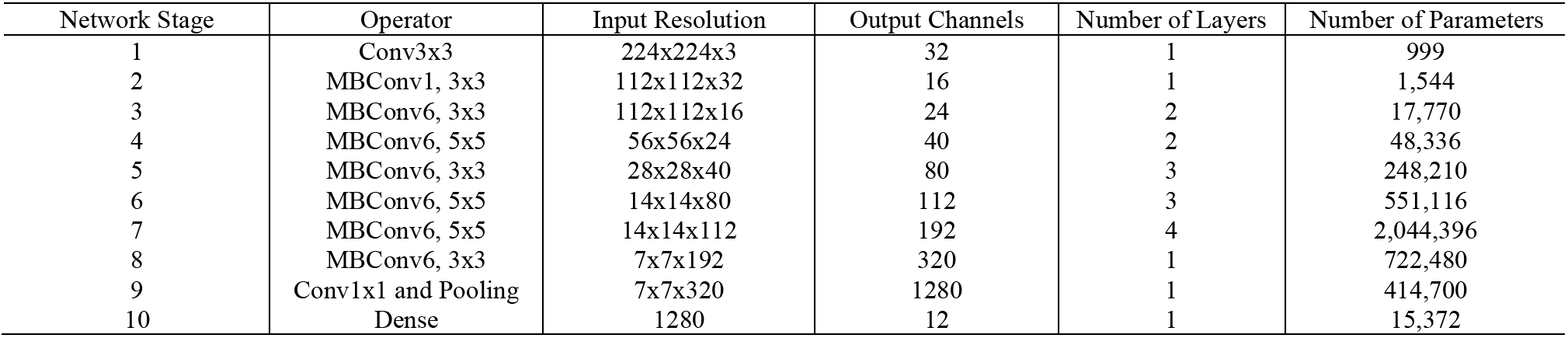
The EfficientNetB0 convolutional neural network used for image classification, including the number of layers, parameters, output channels, and input resolution for each stage. For more information on the architecture design, see [21].

| Network Stage | Operator | Input Resolution | Output Channels | Number of Layers | Number of Parameters |
| --- | --- | --- | --- | --- | --- |
| 1 | Conv3x3 | 224x224x3 | 32 | 1 | 999 |
| 2 | MBConv1, 3x3 | 112x112x32 | 16 | 1 | 1,544 |
| 3 | MBConv6, 3x3 | 112x112x16 | 24 | 2 | 17,770 |
| 4 | MBConv6, 5x5 | 56x56x24 | 40 | 2 | 48,336 |
| 5 | MBConv6, 3x3 | 28x28x40 | 80 | 3 | 248,210 |
| 6 | MBConv6, 5x5 | 14x14x80 | 112 | 3 | 551,116 |
| 7 | MBConv6, 5x5 | 14x14x112 | 192 | 4 | 2,044,396 |
| 8 | MBConv6, 3x3 | 7x7x192 | 320 | 1 | 722,480 |
| 9 | Conv1x1 and Pooling | 7x7x320 | 1280 | 1 | 414,700 |
| 10 | Dense | 1280 | 12 | 1 | 15,372 |

The ExoNet images were split into training (89.5%), validation (3.5%), and testing (7%) sets, the proportions of which are consistent with ImageNet [22]. We experimented with transfer learning of pretrained weights from ImageNet [22] but found no additional performance benefit. Dropout regularization was applied before the final dense layer to prevent overfitting during training such that the network weights were randomly dropped (i.e., activations set to zero) at a rate of 0.5. Images were also randomly flipped horizontally during training to increase stochasticity and promote generalization. We trained the network for 40 epochs using a batch size and initial learning rate of 128 and 0.001, respectively; these hyperparameters were experimentally tuned to maximize performance on the validation set (Fig. 3). The learning rate was reduced during training using a cosine weight decay schedule. We calculated the sparse categorical cross-entropy loss between the labelled and predicted classes and used the Adam optimizer [23], which computes gradients using momentum and an adaptive learning rate, to update the network weights and minimize the loss function. During testing, we used a single central crop of 224×224. Training and inference were performed on a Tensor Processing Unit (TPU) version 3-8 by Google Cloud; these customized chips can allow for accelerated CNN computations with less power consumption.

**Fig. 3.**
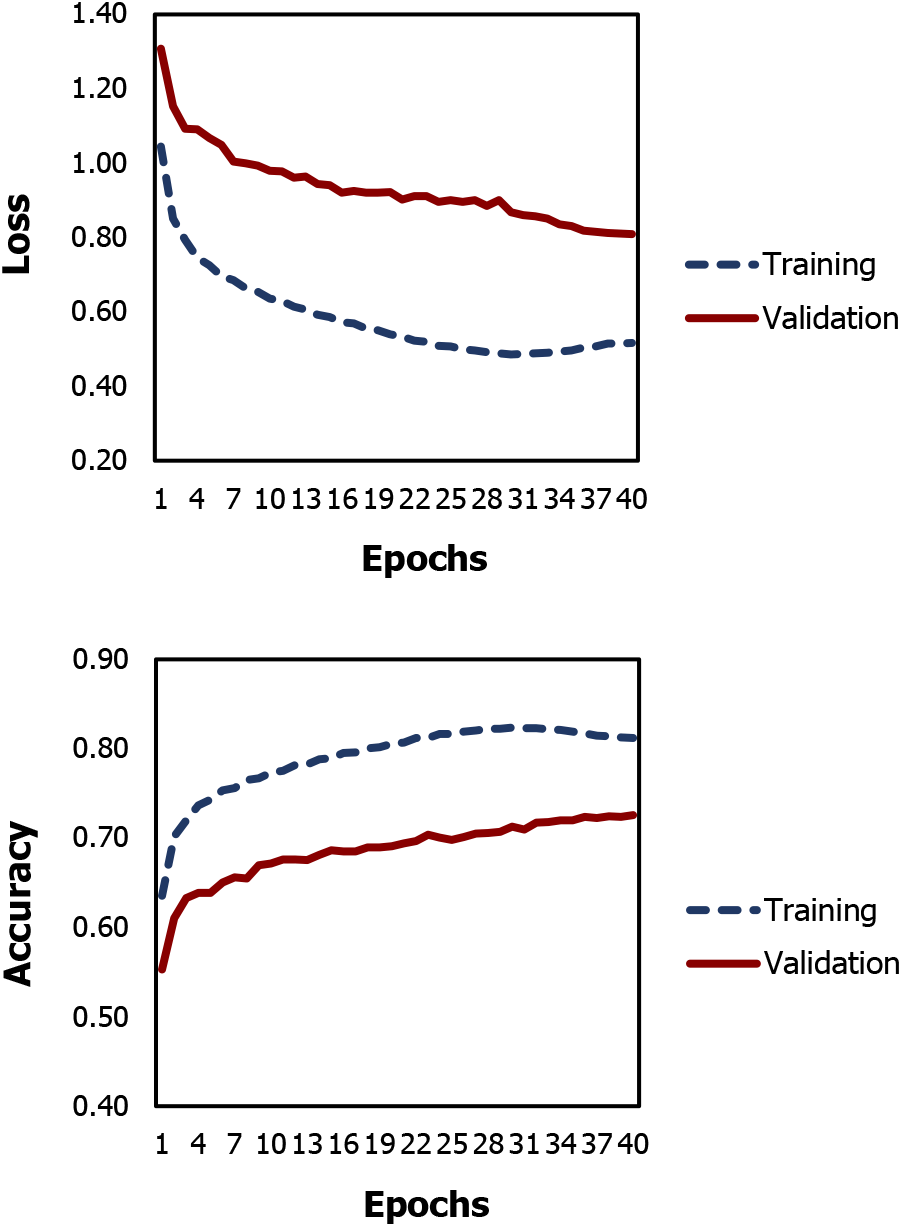
The loss and image classification accuracies during training and validation on the ExoNet database.

## III. Results

The image classification accuracies on the training and validation sets were ~81.2% and ~72.6%, respectively. Table 2 presents the multiclass confusion matrix, which visually illustrates the CNN classification performance during inference. The matrix columns and rows are the predicted and labelled classes, respectively. The diagonal elements are the individual classification accuracies for each environment class, known as true positives, and the nondiagonal elements are the misclassification percentages. Our environment recognition system achieved ~73.2% classification accuracy on the testing set, that being the percentage of true positives (i.e., 47,265 images) from the total number of images (i.e., 64,568 images).

**Table 2.**
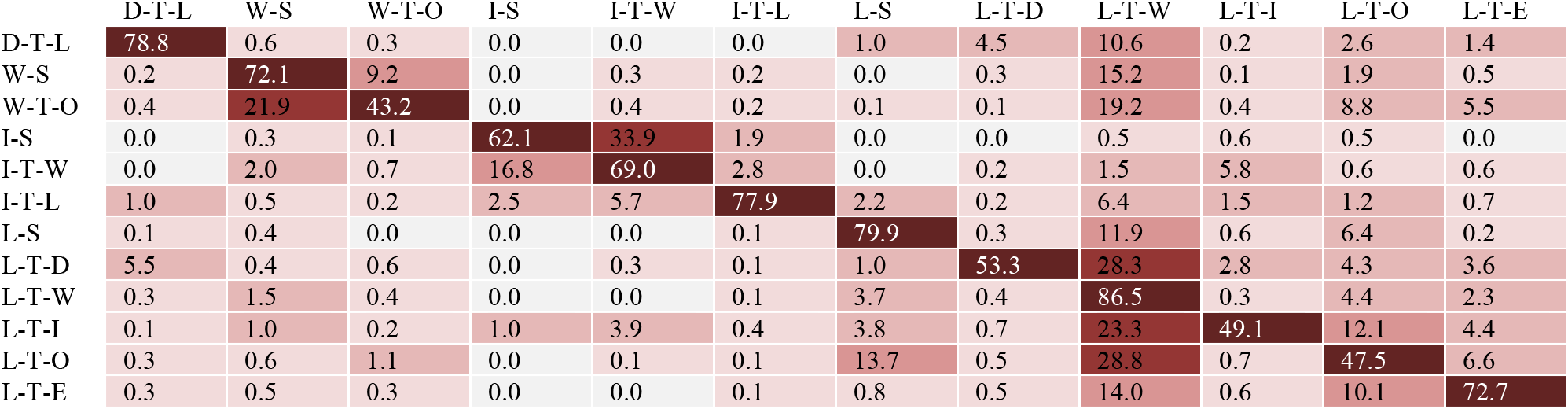
The multiclass confusion matrix illustrating the image classification performance for each environment class during inference. The matrix columns and rows are the predicted and labelled classes, respectively.

|  | D-T-L | W-S | W-T-O | I-S | I-T-W | I-T-L | L-S | L-T-D | L-T-W | L-T-I | L-T-O | L-T-E |
| --- | --- | --- | --- | --- | --- | --- | --- | --- | --- | --- | --- | --- |
| D-T-L | 78.8 | 0.6 | 0.3 | 0.0 | 0.0 | 0.0 | 1.0 | 4.5 | 10.6 | 0.2 | 2.6 | 1.4 |
| W-S | 0.2 | 72.1 | 9.2 | 0.0 | 0.3 | 0.2 | 0.0 | 0.3 | 15.2 | 0.1 | 1.9 | 0.5 |
| W-T-O | 0.4 | 21.9 | 43.2 | 0.0 | 0.4 | 0.2 | 0.1 | 0.1 | 19.2 | 0.4 | 8.8 | 5.5 |
| I-S | 0.0 | 0.3 | 0.1 | 62.1 | 33.9 | 1.9 | 0.0 | 0.0 | 0.5 | 0.6 | 0.5 | 0.0 |
| I-T-W | 0.0 | 2.0 | 0.7 | 16.8 | 69.0 | 2.8 | 0.0 | 0.2 | 1.5 | 5.8 | 0.6 | 0.6 |
| I-T-L | 1.0 | 0.5 | 0.2 | 2.5 | 5.7 | 77.9 | 2.2 | 0.2 | 6.4 | 1.5 | 1.2 | 0.7 |
| L-S | 0.1 | 0.4 | 0.0 | 0.0 | 0.0 | 0.1 | 79.9 | 0.3 | 11.9 | 0.6 | 6.4 | 0.2 |
| L-T-D | 5.5 | 0.4 | 0.6 | 0.0 | 0.3 | 0.1 | 1.0 | 53.3 | 28.3 | 2.8 | 4.3 | 3.6 |
| L-T-W | 0.3 | 1.5 | 0.4 | 0.0 | 0.0 | 0.1 | 3.7 | 0.4 | 86.5 | 0.3 | 4.4 | 2.3 |
| L-T-I | 0.1 | 1.0 | 0.2 | 1.0 | 3.9 | 0.4 | 3.8 | 0.7 | 23.3 | 49.1 | 12.1 | 4.4 |
| L-T-O | 0.3 | 0.6 | 1.1 | 0.0 | 0.1 | 0.1 | 13.7 | 0.5 | 28.8 | 0.7 | 47.5 | 6.6 |
| L-T-E | 0.3 | 0.5 | 0.3 | 0.0 | 0.0 | 0.1 | 0.8 | 0.5 | 14.0 | 0.6 | 10.1 | 72.7 |

The network most accurately predicted the “level-ground transition wall/door” (L-T-W) class with ~86.5% accuracy, followed by “level-ground steady” (L-S) at ~79.9% and “decline stairs transition level-ground” (D-T-L) at ~78.8%. These results could be attributed to the class imbalances among the training data (i.e., there were significantly more images of L-T-W environments compared to other classes). However, some classes with limited images showed relatively good classification performance. For instance, the “incline stairs transition level-ground” (I-T-L) class comprised only ~1.2% of the ExoNet database but had ~77.9% classification accuracy. The least accurate predictions were for the environment classes with *other* features – “wall/door transition other” (W-T-O) at ~43.2% and “level-ground transition other” (L-T-O) at ~47.5%. The average inference runtime was ~2.5 ms/image on the Cloud TPU using a batch size of 8.

## IV. Discussion

Inspired by the human vision-locomotor control system, computer vision can provide important environmental context and features for robotic exoskeleton control. However, small-scale and private training datasets have hindered the development of image classification algorithms for environment recognition [20]. To address these limitations, we developed ExoNet - the largest and most diverse open-source dataset of wearable camera images of walking environments [11]. Unparalleled in both scale and diversity, ExoNet contains over 5.6 million images of indoor and outdoor real-world environments, of which ~923,000 images were annotated using a 12-class hierarchical labelling architecture; these design features are important since deep learning requires significant and diverse training data. We then trained and tested the Efficient-NetB0 convolutional neural network [21] on the ExoNet database to predict the different walking environments, therein providing a benchmark performance for future comparisons. We chose EfficientNetB0 because the architecture design was optimized for both classification accuracy and computational complexity, the features of which are pertinent to onboard real-time inference for robotic exoskeleton control.

Our preliminary environment recognition system achieved ~73% classification accuracy on ExoNet. However, for environment-adaptive control of robotic exoskeletons, near perfect classification accuracy is desired since even rare misclassifications can cause loss-of-balance and injury [24]. Future work could use temporal information to improve the classification accuracy and robustness. Sequential images could be classified using majority voting [5], [16]–[17] or deep learning models like transformers or recurrent neural networks (RNNs) [18]. RNNs process sequential inputs while maintaining an internal hidden state vector that implicitly contains temporal information. However, training RNNs can be challenging due to exploding and vanishing gradients [25]. While these networks were designed to learn long-term dependencies, research has shown that they struggle with storing sequential information over long periods [25]. To mitigate this issue, RNNs can be supplemented with explicit memory modules like long short-term memory (LSTM) networks. A recent study [18] showed that fusing sequential decisions using recurrent neural networks or LSTM networks significantly outperformed CNNs alone for image classification of walking environments. However, using temporal information for environment classification can lead to longer decision times and therefore impede real-time exoskeleton control.

A potential limitation of the ExoNet database is the 2D nature of the environment information. Whereas an RGB camera like ours measures light intensity information [8]–[13], depth cameras can also provide distance measurements [14]–[19]. Depth cameras work by emitting infrared light and calculating distances by measuring the time-of-flight between the camera and surrounding environment. One advantage of depth sensing is the ability to extract environmental features like step height and width, which can improve the mid-level exoskeleton control. However, depth measurement accuracy typically degrades in outdoor lighting conditions and with increasing distance [16]–[17]. Most environment recognition systems using depth cameras have been tested in controlled indoor environments and/or have had limited capture volumes (i.e., 1-2 m of maximum range imaging) [14]–[17]. Moreover, the application of depth cameras for environment sensing would require robotic exoskeletons to have onboard microcontrollers with high computing power and low power consumption; the current embedded systems would need significant modifications to support real-time processing of depth images [16]. These limitations motivated our decision to use RGB images.

Note that since the environmental context does not explicitly represent the user’s locomotor intent, data from computer vision should supplement, rather than replace, the locomotion mode control decisions based on information from surface EMG and/or mechanical and inertial sensors. For instance, images from our wearable smartphone camera system could be fused with the onboard IMU measurements for high-level exoskeleton control. Suppose an exoskeleton user unexpectedly stops before ascending an incline staircase; the acceleration data would indicate static standing rather than stair ascent, despite the staircase being detected within the field-of-view. The onboard IMU measurements could also be used to control the camera’s sampling rate [7]–[8]. Whereas fast walking can benefit from higher sampling rates for continuous classification, standing still does not necessarily require environment information and thus the camera could be powered down, or the sampling rate decreased, to lessen the computational and memory storage requirements. The optimal method for fusing the acceleration data with images for environment-adaptive control of robotic exoskeletons remains to be determined.

## Acknowledgment

We gratefully recognize the TensorFlow Research Cloud Program by Google and the NVIDIA GPU Grant Program for providing the deep learning hardware.

